# Learning to blink strategically is crucial to performance in a predictable saccade task and varies across the lifespan

**DOI:** 10.1101/2025.08.25.670512

**Authors:** Isabell C. Pitigoi, Donald C. Brien, Heidi C. Riek, Blake K. Noyes, Rachel Yep, Olivia G. Calancie, Ryan H. Kirkpatrick, Brian C. Coe, Michele Morningstar, Douglas P. Munoz

## Abstract

Humans blink their eyes 16-20 times each minute to spread tear film on the cornea, representing a substantial amount of waking time when one’s eyes are closed. These spontaneous blinks are strategically timed to prioritize the processing of important visual input, balancing both stimulus characteristics and personal goals. Until now, the learning process underlying optimal blink timing has not been investigated in detail. Here, we present video-based eye-tracking data from 703 healthy participants (aged 5-91 years, 470 female) performing a structured interleaved pro-/anti-saccade task, in which we previously found that blink suppression occurs in anticipation of visual stimulus appearance (Pitigoi et al., 2024). Our goals are to understand (1) how participants modify their blink timing according to the temporal contingencies of the task; (2) whether the capacity to optimize blink timing impacts performance; and (3) whether this pattern varies with age. We found that participants quickly and strategically modified their blink distribution to optimize task performance. Blink probability decreased in periods that would compromise anti-saccade execution and increased when visual input was less critical. We also found significant differences in blink patterns and adaptive ability based on cognitive control capacity (indicated by participants’ anti-saccade error rates). Furthermore, we demonstrated that blink optimization improves gradually from childhood to early adulthood, before declining with advanced age. This supports a possible link between regulation of blink behavior and age-related changes in learning capacity and inhibitory control across the lifespan.

## Introduction

A growing body of work has demonstrated that blinks are strategically timed to optimize visual processing by minimizing co-occurrence with task-relevant or interesting stimuli (Hoppe et al., 2018; Nakano & Miyazaki, 2019; Shultz et al., 2011). Blinks occur before and after periods of information processing in a manner modulated by both sensory signals and cognitive goals (Murali & Händel, 2021; Siegle et al., 2008; Wascher et al., 2015). Thus, this behavior is dependent on and scales with a task’s temporal predictability (Brych & Händel, 2020; Huber et al., 2022, 2023). Furthermore, correlations between total blink rate and task performance outcomes were weak (Pitigoi et al., 2024) or non-existent (Hoppe et al., 2018) on predictable tasks, reinforcing that a higher blink frequency does not itself impair performance—instead, timing matters. Altogether, these findings reinforce that blink behavior is a complex product of both top-down factors related to attention, cognition, and goal-driven behavior, and bottom-up factors related to sensory input. Efficient visual processing requires a controlled balance between these traits which is likely shaped through learning within a task (i.e., to predict the appearance of relevant stimuli), as well as through lifelong experience.

In a study by Hoppe et al. (2018), participants were found to quickly adapt to regularities within the task’s structure and time their blinks strategically according to event probability (i.e., the appearance of visual stimuli). To our knowledge, this is the only study to explore the learning process behind strategic blink behavior. It remains to be determined how quickly strategic blink patterns are established, how they influence performance, and how this varies between individuals. Here we aim to address these questions by analyzing video-based eye-tracking data from a large cohort of healthy participants (n=703) performing the interleaved pro-/anti-saccade task (IPAST). Anti-saccade trials (looking away from a visual stimulus) are more difficult to perform than pro-saccade trials (looking towards a visual stimulus), sometimes producing direction errors due to weak inhibitory control (the ability to suppress automatic responses in favor of appropriate goal-directed behavior) (Coe & Munoz, 2017; Munoz & Everling, 2004). We previously described a consistent pattern for blinking in this task which was driven by its highly predictable stimulus timing structure (Pitigoi et al., 2024). Our first aim is to understand how blink patterns are established and optimized to suit task goals. We expect participants’ blink behavior to quickly change from reactive (i.e., inhibited following stimulus appearance) to predictive (i.e., inhibited in anticipation of stimulus appearance). Previously, we linked anti-saccade direction errors to a failure to suppress blinks during critical periods (Pitigoi et al., 2024). If blink timing is indeed critical for task performance due to shared circuitry related to inhibitory control, we expect blink patterns to be less optimized in participants with high anti-saccade error rates.

While learning to optimize visual behavior may occur over short timeframes (i.e., within a task), it may also improve with accrued experience over longer timeframes (i.e., across the lifespan). Such changes across the lifespan may reflect complex structural and functional changes occurring in the brain to support cognitive processes related to learning capacity and inhibitory control (Ferguson et al., 2021; Luna et al., 2010). The rapid maturation of such processes in adolescence, followed by their gradual decline with aging, brings into question whether the ability to optimize blink timing also varies with age. Our final goal is therefore to address this question by comparing blink behavior across the lifespan; the wide age range of our cohort (5–91 years) permits analysis of healthy development and aging in a single study. We expect the speed and efficiency of blink optimization to improve from childhood to early adulthood, before declining later in life. This trajectory would be complementary to that of anti-saccade error rates and reaction times (Yep et al., 2022) as well as broader visuomotor adaptability across the lifespan (Ruitenberg et al., 2023).

## Materials and Methods

### Participants

All experimental protocols were reviewed for ethical compliance by the Queen’s University Faculty of Health Sciences and Affiliated Teaching Hospitals Research Ethics Board (Protocol #6040314, previously #6005163). We recruited healthy males and females aged 5 and older from the greater area of Kingston, Ontario, Canada who had normal or corrected-to-normal vision and no self-reported history of major neurological or psychiatric illness. The present study includes the same participants as our previous work with the same task (Yep et al., 2022), as well as an additional 140 participants recruited since 2022, for a total of 771. After exclusions (see Table 1), 703 participants (470 female, aged 5-91 years) remained. As evidenced by the large effect sizes in our previous work, this sample size is ample to detect meaningful differences in blink probability between groups (Pitigoi et al., 2024). Participant demographic characteristics are summarized in Table 2. Written informed consent/assent was provided by all participants prior to the start of the experiment. Recording sessions lasted approximately one hour and participants were remunerated $20 CAD for their time. Prior to administration of the task, participants aged ≥18 completed a Montreal Cognitive Assessment (MoCA): a brief screening tool developed to detect mild cognitive impairment (Nasreddine et al., 2005). We reported mean MoCA score for each subgroup of our cohort in Table 2. No exclusions were made based on MoCA score in order to capture a wide range of cognitive ability in our data.

**Table 1.** Reasons for excluding each participant.

| Reason | Num. Participants Excluded |
| --- | --- |
| File corrupted/missing | 1 |
| Calibration failed | 15 |
| >40% of data contained eye loss | 4 |
| Average validation error of more than 1.5° | 12 |
| Less than 240 trials completed | 32 |

**Table 2.** Summary of demographic information for included participants.

| Analysis | Bin (n) | Mean Age<br>± SD | Sex (n) | Mean MOCA<br>± SD | Mean<br>Error Rate<br>± SD |
| --- | --- | --- | --- | --- | --- |
| **Cohort** | Total (703) | 31.0 ± 21.6 | F (470);<br>M (233) | 27.8 ± 2.0<br>(320 missing) | — |
| **Error**<br>**Rate** | <0.075 (97) | 31.8 ± 17.4 | F (65);<br>M (32) | 28.3 ± 1.4<br>(38 missing) | — |
|  | 0.075-0.12 (99) | 29.9 ± 17.6 | F (67);<br>M (32) | 28.0 ± 2.0<br>(38 missing) | — |
|  | 0.12-0.16 (98) | 36.0 ± 21.7 | F (59);<br>M (39) | 27.8 ± 2.2<br>(31 missing) | — |
|  | 0.16-0.235 (99) | 32.3 ± 21.1 | F (68);<br>M (31) | 27.8 ± 2.0<br>(40 missing) | — |
|  | 0.235-0.31<br>(104) | 28.8 ± 20.8 | F (80);<br>M (24) | 27.9 ± 1.5<br>(49 missing) | — |
|  | 0.31-0.45 (105) | 30.4 ± 25.0 | F (73);<br>M (32) | 27.6 ± 1.7<br>(58 missing) | — |
|  | >0.45 (101) | 27.88 ± 25.7 | F (58);<br>M (43) | 26.1 ± 2.9<br>(66 missing) | — |
| **Age** | **YC: Under 10**<br>**(31)** | 8.2 ± 1.2 | F (14)<br>M (17) | — | 0.5421 |
|  | 10-12 (57) | 11.8 ± 0.9 | F (27)<br>M (30) | — | 0.4046 |
|  | 13-14 (94) | 14.0 ± 0.6 | F (66)<br>M (28) | — | 0.3004 |
|  | 15-17 (64) | 16.2 ± 0.9 | F (46)<br>M (18) | — | 0.2135 |
|  | 18-20 (92) | 19.6 ± 0.9 | F (66)<br>M (26) | 28.0 ± 1.6<br>(21 missing) | 0.1974 |
|  | **EA: 21-24 (99)** | 22.4 ± 1.2 | F (69)<br>M (30) | 28.0 ± 1.7<br>(37 missing) | 0.1674 |
|  | 25-34 (54) | 28.9 ± 2.9 | F (37)<br>M (17) | 28.3 ± 1.4<br>(5 missing) | 0.1700 |
|  | 35-44 (37) | 41.2 ± 2.6 | F (30)<br>M (7) | 28.2 ± 1.7<br>(4 missing) | 0.1339 |
|  | 45-54 (41) | 50.0 ± 2.8 | F (28)<br>M (13) | 28.3 ± 1.5<br>(1 missing) | 0.2146 |
|  | 55-70 (76) | 63.0 ± 4.5 | F (51)<br>M (25) | 27.7 ± 1.8<br>(7 missing) | 0.2202 |
|  | **OA: 70 and**<br>**over (58)** | 78.2 ± 6.1 | F (36)<br>M (22) | 26.0 ± 2.9<br>(4 missing) | 0.3185 |

118 MOCA = Montreal Cognitive Assessment Score, F = female, M = male.

### Eye-Tracking Apparatus

A video-based eye tracker (Eyelink-1000 Plus monocular-arm; SR Research Ltd, ON, Canada) was used to record right eye position and pupil area monocularly at 500 Hz. In cases where right eye calibration was difficult, the left eye was tracked instead. Participants were seated 60cm from the screen (a 17-inch LCD monitor controlled through a Dell Latitude E7440 Laptop; 1280×1024 pixels, 32-bit color, 60Hz refresh rate) in a dark, windowless room. Gaze position was calibrated and validated using a 9-point array (in 63 participants, due to difficulties calibrating, a 5-point array was used instead). An average validation error of less than 1° was considered sufficient for inclusion in analyses.

### Task

Participants completed the interleaved pro-/anti-saccade task (IPAST; Fig. 1) in two blocks of 120 trials, each approximately 7 min in duration. Every 40 trials, a brief calibration check took place which verified the accuracy of eye-tracking and permitted re-calibration if necessary. Every trial began with the appearance of a central fixation cue (0.5° diameter circle, 42cd/m^2^; 1000 ms duration), the color of which indicated the saccade instruction (green = pro-saccade, red = anti-saccade). This fixation epoch (FIX) was followed by a GAP period lasting 200 ms in which the screen was completely black, after which a peripheral target stimulus (STIM; 0.5° diameter circle; white, 62cd/m^2^; 1000 ms duration) appeared 10° to the left or right of screen center. Participants were required to look towards the STIM on pro-saccade trials (automatic response) and opposite to the STIM on anti-saccade trials (voluntary response requiring inhibitory control). Each trial was separated by an intertrial interval (ITI) consisting of a black screen that lasted for 1000 ms. Both saccade instructions (pro-saccade vs. anti-saccade) and STIM locations (left vs. right) were pseudo-randomly interleaved.

**Figure 1.**
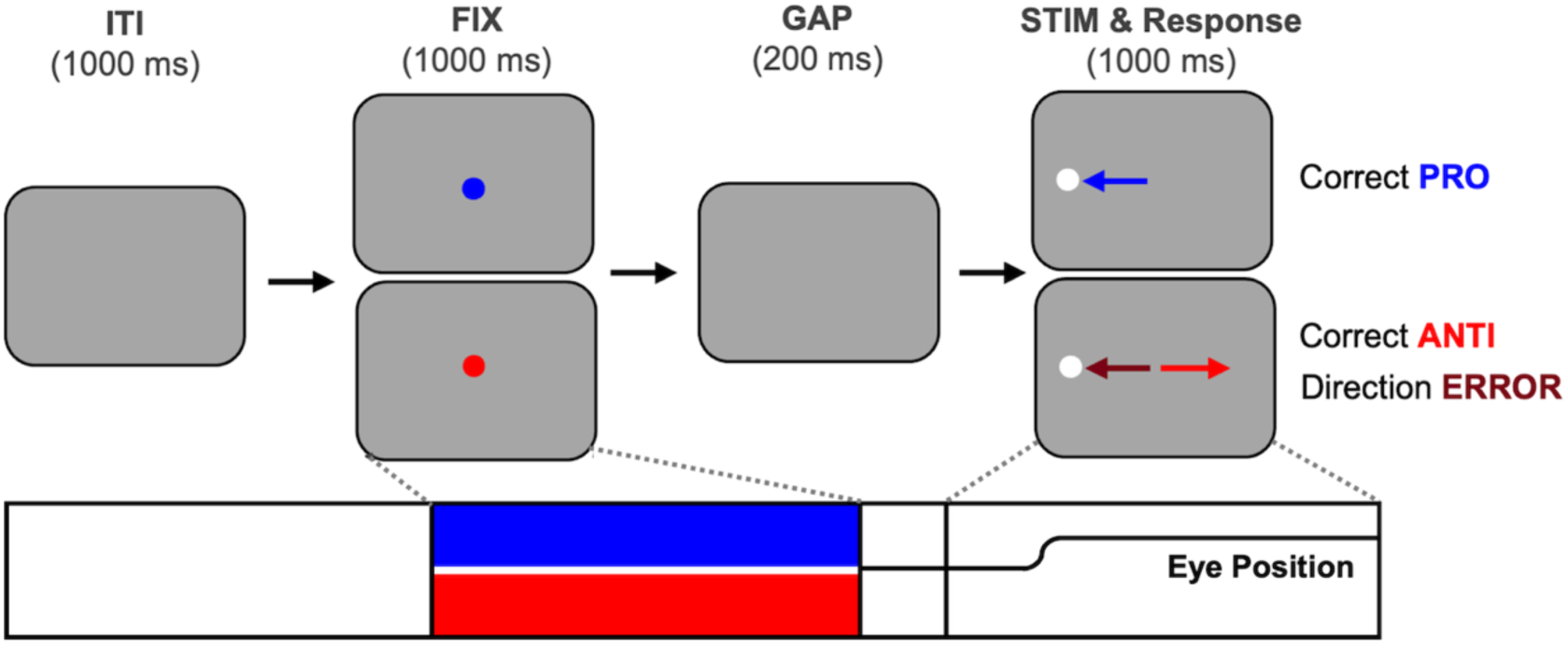
The Interleaved Pro-/Anti-Saccade Task (IPAST). Each trial started with a central fixation cue (FIX), the color of which indicated the trial instruction (pro-saccade = green; anti-saccade = red). Note that pro-saccades are coded blue in this paper. After a 200ms GAP consisting of a blank black screen, a white peripheral stimulus (STIM) appeared at 10° horizontally to the left or right of where FIX had been. After STIM onset, the eye position trace represents a saccade being made in the appropriate epoch.

### Statistical Analyses

All data analysis was performed using custom programs built in MATLAB (MathWorks Inc, Natwick, MA, USA).

#### Pre-Processing

All data were pre-processed using in-house auto-marking code for blink and saccade detection (Coe et al., 2024). Trials were categorized as correct vs. incorrect depending on the direction of the first saccade after STIM onset. Blink start and end times were defined by modelling the noise in pupil area data which flanks the periods of data loss caused by blinks. In our previous work (Pitigoi et al., 2024), we defined viable blinks as those lasting 50 to 500 ms. Data losses with durations outside of this range were excluded as they were likely tracking artifacts or reflective of participants closing their eyes for other reasons (e.g., fatigue).

Participants were excluded if they had insufficient data to adequately characterize task performance (>40% of data contained eye loss) either due to poor eye-tracking or inability or unwillingness to complete the task. Only participants who completed all 240 trials were included due to the emphasis on analysis spanning the entire length of the task. We also excluded participants with <30 viable trials of each kind (pro- and anti-saccade). Viable trials were defined as having a saccade take place between 90 and 800 ms after STIM onset (Coe et al., 2024).

#### Blink Probability

Blink probability was used to quantify blink behavior as it provides temporal sensitivity and acts as a two-dimensional measure incorporating both rate and duration. A logical array identified when the eye was open (0) or closed (1) in each trial. An average of this array was calculated across all trials for each participant to yield blink probability aligned on time of STIM onset. Correct anti-saccade trials and direction error trials were then averaged separately and compared for significant differences (see *Group Curve Comparisons*) to determine time periods where blinking impacted trial performance. We also plotted blink probability curves on select trials (e.g. 1^st^, 2^nd^, 3^rd^) to capture changes brought on by initial exposure to the task, and in groups of 40 trials (1-40, 41-80, 81-120, 121-160, 161-200, 201-240) to visualize changes across the entire duration of the task.

#### Group Curve Comparisons

To compare time-series blink probability curves between groups, we implemented a statistical method similar to that developed in Pitigoi et al. (2024). Briefly, we fit cubic smoothing splines to the raw participant data (csaps function in MATLAB; smoothing parameter = 0.00001), then used bootstrapping to estimate the mean and 95% confidence intervals (CIs; type = student) by resampling randomly with replacement (1000 iterations). Time points where the 95% CI of the difference of resampled means did not include 0 were recorded. To filter out spurious differences driven by multiple comparisons and noise in the data, permutation testing (1000 iterations) was performed on unlabelled data without replacement to generate a null distribution. For each permutation, we stored the duration of the maximum period of significant difference (representing the longest duration of a false positive result). After 1000 iterations, we calculated the upper 95% CI to establish the threshold for significance by which to filter each comparison (reported in Table 3). Statistical significance was then marked in the real dataset where the curves being compared were significantly different for *at least* that threshold duration of milliseconds consecutively. We have also shown ‘trending’ periods which fall short of the significance cut-off but represent the upper 90% CI, in acknowledgement of this method’s sensitivity to extreme values. Cohen’s d effect sizes and their 95% CIs were also plotted across the length of the trial for each comparison. Cohen’s d values below 0.2 were shown in a lighter color as they indicate a very small effect size and usually do not overlap with significant timepoints.

**Table 3.** Thresholds for statistical significance obtained from permutation analysis.

| Figure | Comparison | Significance threshold (ms) | Trending threshold (ms) |
| --- | --- | --- | --- |
| 2A | Pro-saccade to Anti-saccade | 124 | 94 |
|  | Anti-saccade to Direction Error | 132 | 102 |
|  | Pro-saccade to Direction Error | 134 | 104 |
| 4A | HI error rate to LO error rate | 194 | 150 |
| 5A | YC to EA | 289 | 222 |
| 5B | EA to OA | 228 | 174 |

Thresholds are rounded up to the nearest even number. They indicate the minimum duration for which two curves must differ to establish a significant (upper 95% CI) or trending (upper 90% CI) effect. HI = Highest, LO = Lowest, YC = young children; under 10 years, EA = early adults; 21-24 years, OA = oldest adults; 70 years and older.

#### Learning Trajectories

To understand the learning trajectories of each group, we plotted changes in blink probability occurring in three critical periods: the middle of the ITI (ITI_m_), the middle of FIX (FIX_m_), and the transitionary period between them (TRS_m_). For each trial, we calculated the average blink probability in the 100 ms period representing the middle of each epoch, indicated by the subscript letter m (ITI_m_: −1900 to −1800 ms; TRS_m_: −1250 to −1150 ms; FIX_m_: −800 to −700 ms, relative to STIM onset). Note that the TRS_m_ period captures FIX onset at −1200 ms, and thus represents the transitionary period between ITI and FIX rather than being a distinct epoch in the task. This yielded three arrays (one for each epoch) containing an average value for each of 240 trials for all participants which completed the task in its entirety (n=699, 4 additional participants were excluded at this stage for incompleteness). We fit cubic smoothing splines to each participants’ data across all 240 trials (csaps function in MATLAB; smoothing parameter = 0.0001), then used bootstrapping to estimate the mean and 95% CIs by resampling randomly with replacement (1000 times). To determine where the slope of the mean was changing significantly (i.e. learning was occurring), we calculated its first derivative. Periods of significant change were marked at time points when the 95% CI of the first derivative did not contain 0.

#### Anti-Saccade Error Rate Analysis

Participants’ error rates were calculated by dividing the number of anti-saccade direction error trials by the total number of viable anti-saccade trials. For the previously defined critical periods (ITI_m_, TRS_m_, FIX_m_), as well as equal length periods in the middle of the GAP (−150 to −50 ms) and at STIM onset (−50 to 50 ms), we ran Spearman correlations to investigate associations between participants’ error rates and their average blink probability in that period.

To further examine whether blink patterns are functionally related to anti-saccade performance, we compared blink distribution on anti-saccade trials between participants at different performance levels. Participants were separated into seven tiers based on their error rate such that groups were comparable in size (~100 participants per group; Table 2). We then plotted blink probability relative to STIM onset on anti-saccade trials for all seven groups separately and compared the lowest (LO: <0.075) and highest (HI: >0.45) error rate groups for significant differences (see *Group Curve Comparisons*). Time periods where blink probability was significantly higher in the group with the HI error rate would indicate high risk to task performance, while periods with significantly higher blink probability in the LO error rate group would indicate low risk. The learning trajectories of HI and LO error groups in ITI_m_, TRS_m_, and FIX_m_ were also analyzed described in *Trial-by-Trial Analysis*.

#### Age-Related Changes

To characterize changes in blink probability across the lifespan, we separated participants into 10 groups across the lifespan (age range 5-91 years). Blink probability curves for participants aged 5 to 21 years were plotted separate from those aged 24 and 91 years to examine developmental changes in youth separately from aging in older adulthood. The group aged 21-24 years (early adults; EA) was included in both plots as a point of comparison, as it lies at the intersection between developmental and aging trajectories. We expect blink optimization to peak in this age group, as this was the case for cognitively linked saccade variables in the IPAST (Yep et al., 2022). Age groups were chosen such that those on the same plot were of comparable bin width: youth groups had age ranges of ~2-3 years, while adult groups had age ranges of ~10 years (Table 2). We tested both the youngest children (YC; under 10 years old) and the oldest adults (OA; 70 years and older) against EA for significant differences (see *Group Curve Comparisons*). The learning trajectories of YC, EA, and OA groups in ITI_m_, TRS_m_, and FIX_m_ were also analyzed as in *Trial-by-Trial Analysis*.

## Results

### IPAST Blink Pattern

Our previous work in the IPAST has revealed that blink probability was suppressed in anticipation of FIX and STIM appearance (Pitigoi et al., 2024). The same pattern was found in the expanded cohort of participants in the present study (Fig. 2). Blinks were less likely to occur on correct anti-saccade trials than on direction error trials during FIX onset (−1282 to −940 ms; Cohen’s d ~0.2 to 0.3), as well as from the middle of FIX until just after STIM onset (−624 to 134 ms; Cohen’s d ~0.2 to 1.0), suggesting that blinks at these times pose the highest risk to anti-saccade performance. The effect size for this comparison reached its maximum during the GAP period (Fig. 2B). Blinks were most likely in the ITI across all types of trials, corroborating it as the least risky period to blink in the IPAST independent of trial type.

**Figure 2.**
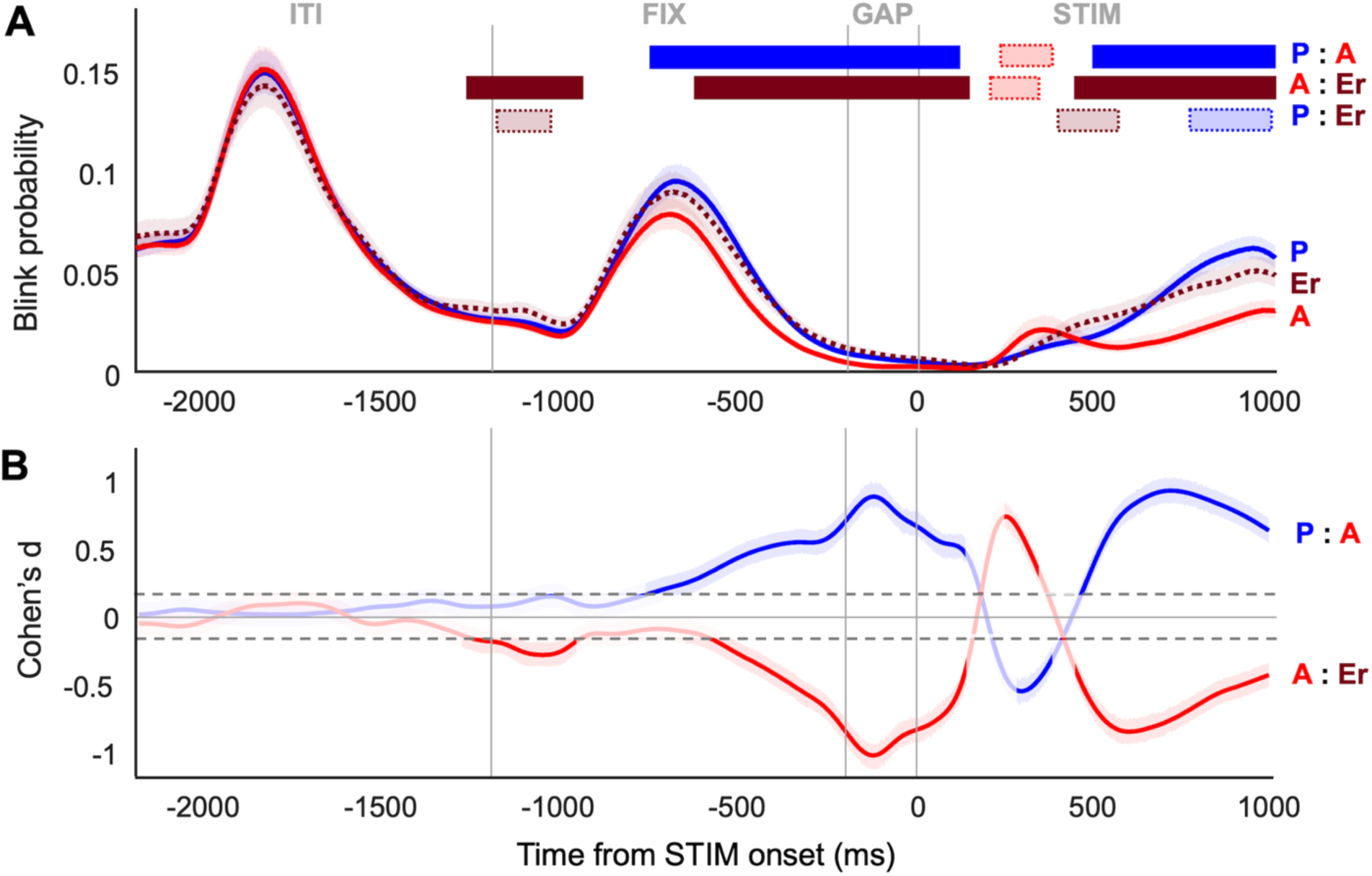
**(A) Blink probability relative to STIM onset, separated by trial type**. Correct pro-saccade = blue; correct anti-saccade = red; direction error = dotted maroon. Shaded area of each curve represents the 95% CI of the bootstrapped means. Horizontal bars along the top of the plot represent periods where differences between the curves are significant (solid color) or trending (pale color with dotted border). Correct pro-saccade trials have a significantly higher blink probability than correct anti-saccade trials from −754 to 104 ms and 494 to 1000 ms. Direction error trials have a significantly higher blink probability than correct anti-saccade trials from −1282 to −940 ms, −624 to 134 ms, and 436 to 1000 ms. **(B) Effect sizes (Cohen’s d) of the mean difference score of blink probabilities for each comparison relative to STIM onset.** Pro-saccade to anti-saccade comparison = blue; anti-saccade to direction error comparison = red. Pro-saccade to direction error comparison not showed due to lack of significant differences. Shaded area of each curve represents the 95% CI of Cohen’s d. Horizontal grey dashed lines represent the threshold for a small effect size (Cohen’s d = ±0.2). Periods that were not significant or did not reach this threshold are shown as faded in color.

### Changes in Blink Behavior Throughout Task

We also explored differences in blink behavior throughout the task as participants learned to optimize their blink timing. We calculated blink probability curves for each trial separately and plotted select trials to highlight changes occurring early in the task (Fig. 3A). In the first two trials (red and orange traces), blinks were suppressed following FIX onset, whereas in the next two trials (yellow and green traces), blinks were suppressed ahead of FIX onset and increased afterwards. This indicated a switch from reaction to anticipation of instruction appearance. From this point onward, this anticipatory pattern became increasingly more pronounced, with blinks increasing in the middle of the ITI and FIX, but not during FIX onset or STIM onset (Fig. 3B).

**Figure 3.**
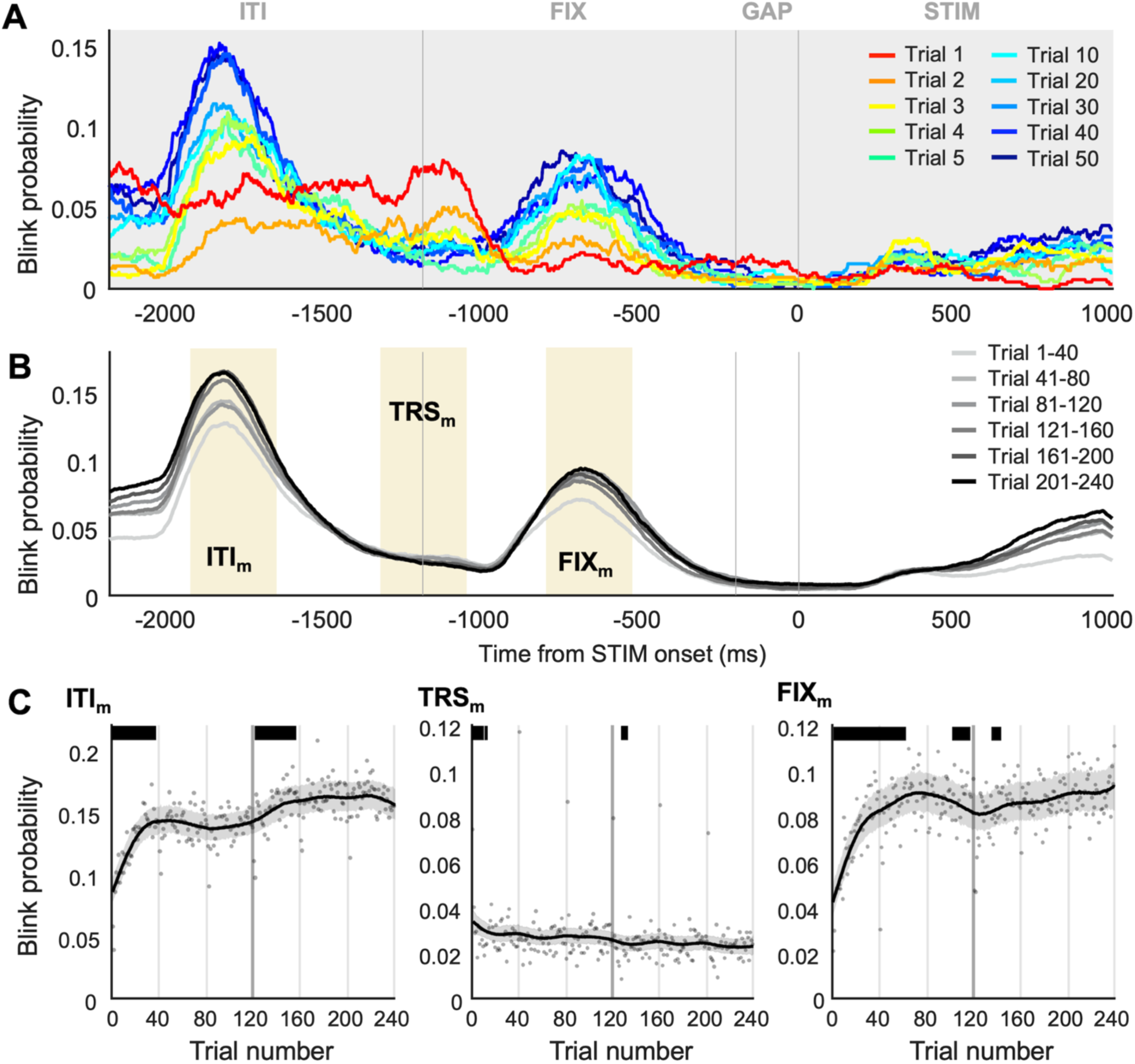
**(A) Blink probability shown relative to STIM onset on selected trials, demonstrating initial adjustments to the task. (B) Blink probability shown relative to STIM onset separated into groups of 40 trials, demonstrating changes across the entire task.** Yellow shading represents the time periods referred to as ITI_m_ (1900 to −1800 ms), TRS_m_ (−1250 to −1150 ms), and FIX_m_ (−800 to −700 ms). **(C) Learning trajectories.** Average blink probability on every trial calculated during ITI_m_ (far left), TRS_m_ (middle), and FIX_m_ (far right). Each point represents an individual trial averaged across all participants. Curves represent the bootstrapped means and shaded areas represent their 95% CIs. Horizontal bars along the top of the plot represent periods where the curve is significantly changing (where the 95% CI of first derivative did not include 0). Vertical gray lines represent the timing of the calibration checks, with the darkest one representing the halfway point of the task (Trial 120).

Analysis of learning trajectories further revealed an increase in blink probability in ITI_m_ during the first ~40 trials, then a brief stabilization followed by another increase at halfway through the task (Fig. 3C, left). During FIX onset (TRS_m_), blink probability appeared to slightly decrease across the entire task, with significant change occurring during the first 15 trials (Fig. 3C, middle). Similar to ITI_m_, blink probability in FIX_m_ increased during the first ~60 trials, then remained relatively stable for the rest of the task aside from a brief dip halfway through the task (Fig. 3C, right). Interestingly, calibration checks were followed by a dramatically lower blink probability on the next trial in ITI_m_ and FIX_m_, and a spike in blink probability in TRS_m_, potentially related to the disruption in task rhythm and participant focus caused by this visual break.

### Relationship to Anti-Saccade Performance

First, we computed correlations to examine the relationship between participants’ blink probability in each epoch and their anti-saccade error rate. Error rate was moderately positively correlated to blink probability during FIX onset (TRS_m_: *rho*=0.3257, *p*<0.001, Spearman), during the GAP (*rho*=0.4440, *p*<0.001, Spearman), and during STIM onset (*rho*=0.4140, *p*<0.001, Spearman). To a lesser degree, greater blink probability during FIX_m_ also corresponded to higher error rate (*rho*=0.1186, *p*=0.0016, Spearman). On the other hand, error rate was weakly negatively correlated to blink probability during ITI_m_ (*rho*=-0.1235, *p*=0.001, Spearman).

Blink probability curves were plotted across a range of error rates in Figure 4. Progressing from the lowest (LO) to the highest (HI) error rate groups, blink probability decreased in the ITI but increased during FIX and STIM onset. The HI error group had significantly higher likelihood to blink during FIX onset (−1584 to −952 ms; Cohen’s d ~0.3 to 0.9) and during STIM onset (−512 to 238 ms; Cohen’s d ~0.3 to 1.7) and lower likelihood to blink in the middle of the ITI (−1990 to −1784 ms; Cohen’s d ~0.2 to 0.6) compared to the LO error group. Figure 4B shows how Cohen’s d values for this comparison fluctuate across the length of a trial, with the effect size peaking near STIM onset. Next, we examined the learning trajectories of HI and LO error groups (Fig. 4C). In the ITI_m_, the LO error group adapted very quickly (within ~30 trials) while the HI error group took longer to stabilize (~ 50 trials). The LO error group also increased their blinks again in the second half of the task while the HI error group did not continue to take advantage of this epoch. In FIX_m_, both groups adapted within the first 30 trials but the trajectory for the LO error group appears to indicate they optimized to a greater degree (i.e., steeper slope, stabilizes at a much higher blink probability despite starting around the same level). Interestingly, in the second half of the task, the two groups are mostly overlapping as blinking in FIX decreases for the LO error group and increases for the HI error group.

**Figure 4.**
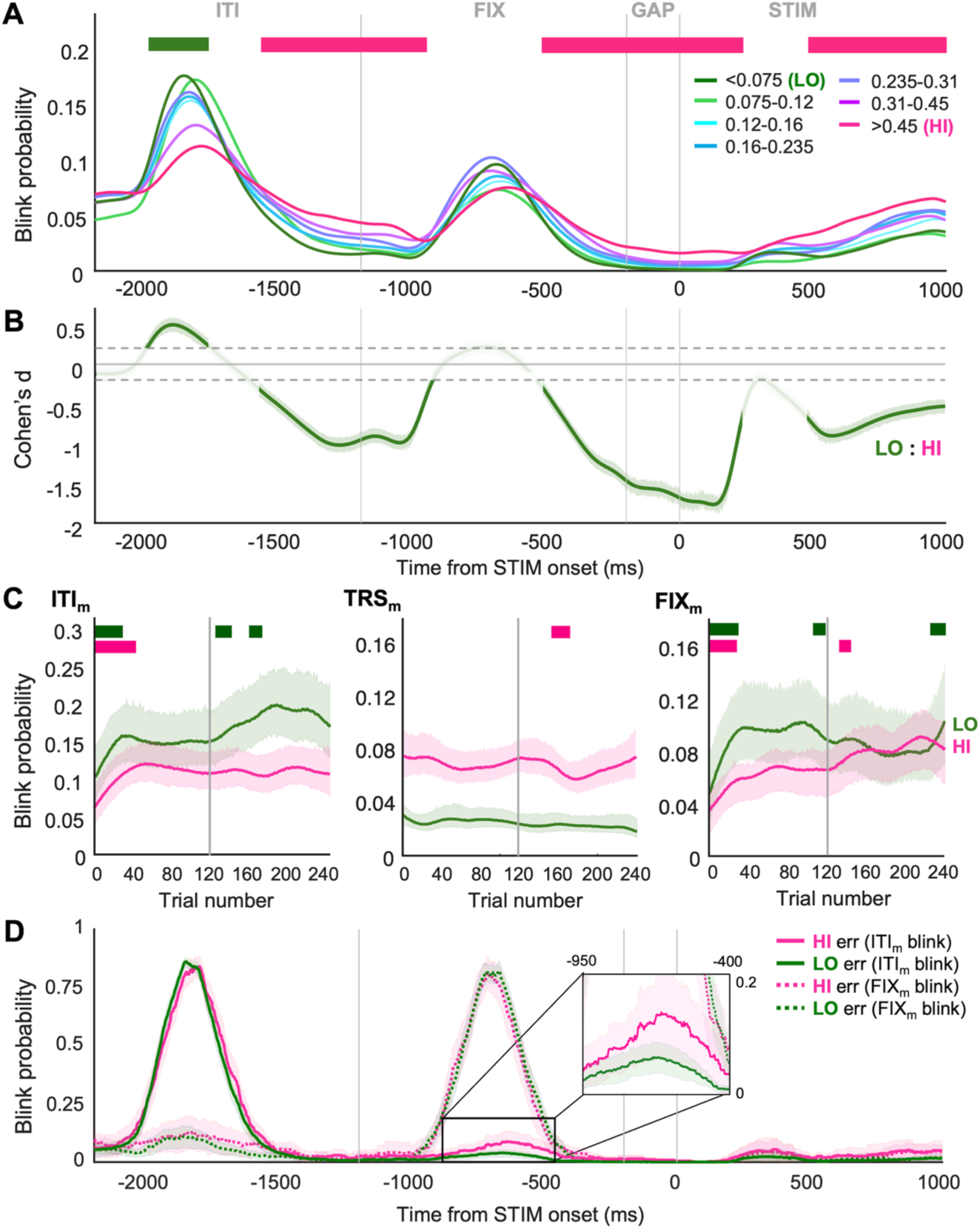
**(A) Blink probability shown relative to STIM onset in participants grouped by their error rate.** Horizontal bars along the top of the plot represent periods where the lowest (LO) and highest (HI) error rate groups differ significantly and are colored according to which group was higher at that time. LO have a significantly higher blink probability than HI from −1990 to −1784 ms, and a significantly lower blink probability than HI from −1584 to −952 ms, −512 to 238, and 472 to 1000 ms. **(B) Effect sizes (Cohen’s d) of the mean difference score of blink probabilities for LO to HI comparison relative to STIM onset.** Shaded area of each curve represents the 95% CI of Cohen’s d. Horizontal grey dashed lines represent the threshold for a small effect size (Cohen’s d = ±0.2). Periods that were not significant or did not reach this threshold are shown as faded. (**C) Learning trajectories.** Average blink probability on every trial calculated during ITI_m_ (far left), TRS_m_ (middle), and FIX_m_ (far right). Each point represents an individual trial averaged across all participants in that group. Curves represent the bootstrapped means and shaded areas represent their 95% CIs. Horizontal bars along the top of the plot represent periods where the curve is significantly changing (where the 95% CI of first derivative did not include 0) and are matched in color to the corresponding group. Vertical gray line represents the calibration check at the halfway point of the task (Trial 120). **(D) Blink probability shown relative to STIM onset for LO and HI error rate groups**, separated based on whether there was a blink in the ITI (solid lines) or in FIX (dashed lines). Shaded area of each curve represents the 95% CIs of the bootstrapped means. Indent zoomed in on area from −950 to −400 ms on x-axis, and 0 to 0.2 on y-axis.

Lastly, we compared trials with blinks in ITI_m_ to those with blinks in FIX_m_, to investigate whether different strategies were more likely to produce additional blinks elsewhere in the trial and thus be less efficient (Fig. 4D). Overall, we found multiple blinks on the same trial to be rare (~14% of all trials). However, in the HI error groups, blinks in the ITI were more often accompanied by an additional blink in FIX (although this did not reach significance).

### Age-Related Changes Across Lifespan

To investigate age-related changes in blink timing across the lifespan, we plotted blink probability in discrete age bins from 5 to 91 years of age (Fig. 5). Within the child and adolescent range (Fig. 5A; ages 5-24 years), each subsequent older age group appeared more efficient in their blink timing, with increasing blink probability during lower-risk periods and decreasing blink probability during higher-risk periods. Statistical analysis of the youngest children (YC; under 10 years) and early adults (EA; 21-24 years) revealed the latter had significantly higher blink probability during the middle of the ITI (−2032 to −1656 ms; Cohen’s d ~0.2 to 0.8), and significantly lower blink probability during FIX onset (−1480 to −990 ms; Cohen’s d ~0.2 to 0.9) and during STIM onset (−432 to 236 ms; Cohen’s d ~0.5 to 1.3). There was an additional period where YC had lower blink probability in the middle of FIX (−930 to −630 ms; Cohen’s d ~0.4 to 0.6), though the duration of this period qualified it as trending and not significant. Figure 5C shows how Cohen’s d values for this comparison fluctuate across the length of a trial, with the largest effect size occurring near STIM onset.

**Figure 5.**
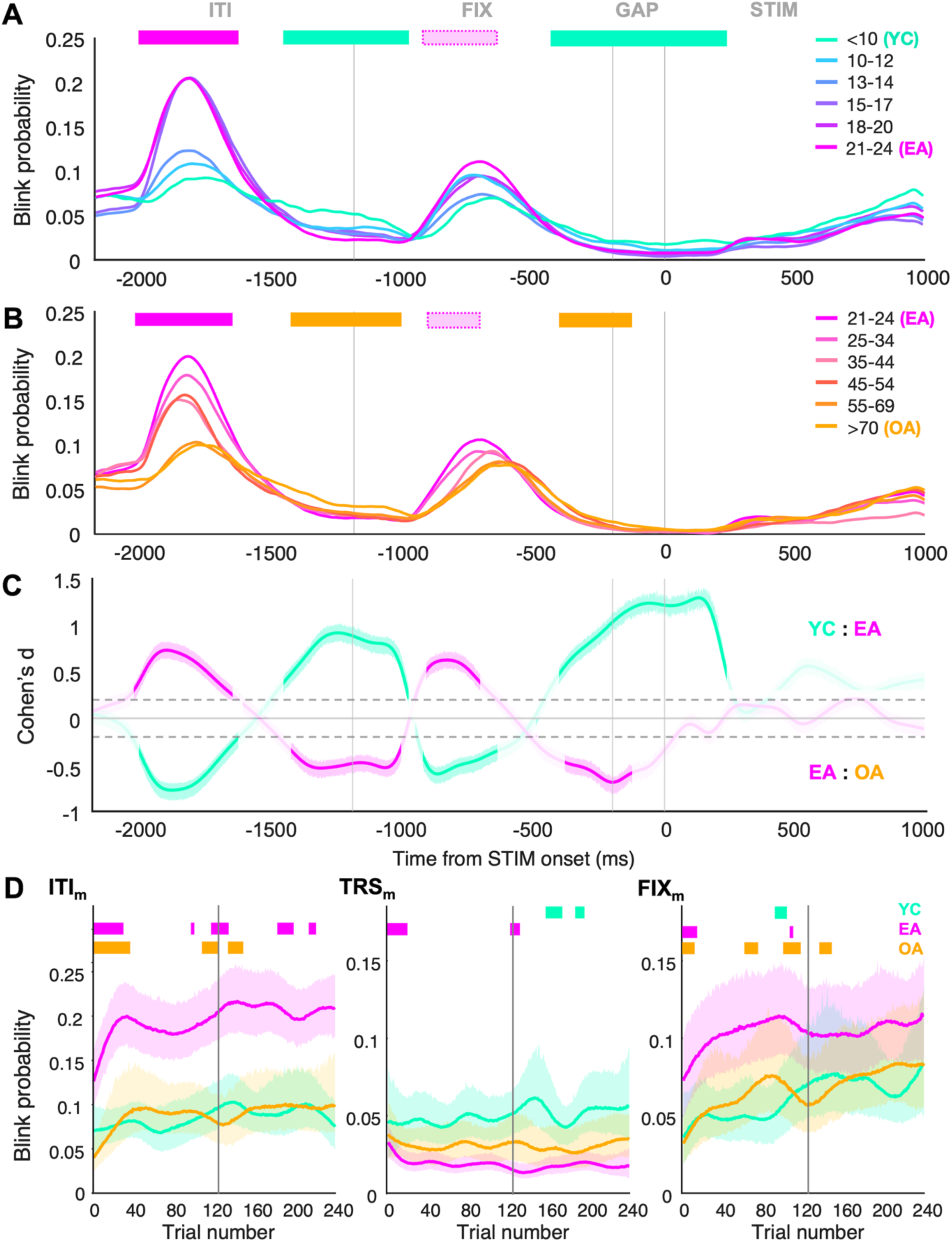
**(A) Blink probability shown relative to STIM onset in participants grouped by age, ranging from 5-24 years.** Horizontal bars along the top of the plot represent periods where differences between YC and EA groups are significant (solid color) or trending (pale color with dotted border) and are colored according to which group was higher at that time. EA have significantly higher blink probability than YC from −2032 to −1656 ms, and significantly lower blink probability than YC from −1480 to −990 ms and −432 to 236 ms. Vertical gray lines represent the boundaries of ITI, FIX, GAP, and STIM epochs indicated along the top of the figure. **(B) Blink probability shown relative to STIM onset in participants grouped by age, ranging from 21-91 years.** Horizontal bars along the top of the plot represent periods where differences between EA and OA groups are significant (solid color) or trending (pale color with dotted border) and are colored according to which group was higher at that time. EA have significantly higher blink probability than OA from −2030 to −1676 ms, and significantly lower blink probability than OA from −1418 to −1016 ms and −398 to −124 ms. **(C) Effect sizes (Cohen’s d) of the mean difference score of blink probabilities for each comparison relative to STIM onset.** YC to EA comparison = teal; EA to OA comparison = magenta. Shaded area of each curve represents the 95% CI of Cohen’s d. Horizontal grey dashed lines represent the threshold for a small effect size (Cohen’s d = ±0.2). Periods that were not significant or did not reach this threshold are shown as faded. **(D) Learning trajectories.** Average blink probability on every trial calculated during ITI_m_ (far left), TRS_m_ (middle), and FIX_m_ (far right). Each point represents an individual trial averaged across all participants in that group. Curves represent the bootstrapped means and shaded areas represent their 95% CIs. Horizontal bars along the top of the plot represent periods where the curve is significantly changing (where the 95% CI of first derivative did not include 0) and are matched in color to the corresponding group. Vertical gray line represents the calibration check at the halfway point of the task (Trial 120).

As expected, the inverse relationship was observed into older adulthood (Fig. 5B; ages 24-91 years), with each subsequent older age group being less strategic in blink timing. Aging was associated with gradually higher likelihood to blink in higher-risk periods and lower likelihood to blink in lower-risk periods. Statistical analysis of EA and oldest adults (OA; 70 years and older) revealed the latter had significantly higher blink probability during FIX onset (−1418 to −1016 ms; Cohen’s d ~0.3 to 0.5) and near the GAP interval (−398 to −124 ms; Cohen’s d ~0.5 to 0.7), as well as significantly lower blink probability in the middle of the ITI (−2030 to −1676 ms; Cohen’s d ~0.2 to 0.7). Again, there was an additional period where OA had lower blink probability in the middle of FIX (−928 to −710 ms; Cohen’s d ~0.3 to 0.6), but the duration of this period qualified it as trending and not significant. Figure 5C shows how Cohen’s d values for this comparison fluctuate across the length of a trial, with the largest effect size occurring in the middle of the ITI and at the start of the GAP interval. The effect was overall weaker than for the comparison of YC and EA. The peak in FIX also qualitatively appeared to be delayed throughout aging.

Last, we plotted the learning trajectories of YC, EA, and OA (Fig. 5D). In ITI_m_ and FIX_m_, EA and OA groups appeared to adapt rapidly to the task, unlike the YC group which did not show any significant change across the task. During FIX onset (TRS_m_), only the EA group showed evidence of adaptation (i.e. a significant decrease in blinks at the beginning of the task). Across all trials, the EA group also maintained the highest blink probability during ITI_m_ and FIX_m_, and the lowest blink probability during TRS_m_, consistent with their behavior being best optimized to high- and low-risk periods in the task.

## Discussion

Here, we investigated how an optimal blink pattern was learned over the course of a structured visual task—a question which has historically not received enough attention. While the literature is rich in evidence of blink patterns being driven by a task’s visual stimuli (Brych & Händel, 2020; Calancie et al., 2022; Nakano et al., 2009; Shultz et al., 2011), the process through which these patterns emerge is rarely studied (Hoppe et al., 2018). Here, we analyzed eye-tracking data from healthy participants performing the IPAST. The predictable timing of stimulus appearance in the IPAST made it ideal for understanding the adaptive nature of blinking. Participants rapidly modified their blinking to limit interference with visual stimuli (instructions, targets) appearing at fixed times in each trial. Blink patterns were most optimal in early adults (21-24 years), as well as in participants with very low anti-saccade error rates. Overall, these findings support a functional role for blink optimization and suggest a link to high-level circuitry involved in learning and regulating behavior on a broader scale.

### Learning of Task-Optimal Blink Timing

Our data demonstrate that, in a task with fixed timing structure, participants quickly learn to blink before stimulus onset rather than during it, indicating rapid adaptation to the task’s timing and risk profile even by the third trial (Fig. 3A). Blink probability increased in the middle of the ITI and FIX epochs over the first ~40 and ~60 trials, respectively, before stabilizing. This echoes prior studies showing that performers of a visual task quickly reached a steady-state for blink behavior which balanced both task-related and physiological costs (Calancie et al., 2022; Hoppe et al., 2018). Beyond the initial learning period, we saw moderate increases in blink probability in the ITI (after Trial 120; Fig. 3C, left). These changes occurring later in the task are likely attributed to participants fatiguing after ~5 minutes of screen time, as both ocular (dry eye, blue light exposure) and mental (sustained attention) strain have been found to increase blink rate on visual tasks after as little as 4 minutes (Maffei & Angrilli, 2019; Stern et al., 1994). Notably, surplus blinks usually occurred in the middle of the ITI, where they would be least disruptive, and not in high-risk periods like during FIX onset. Conceivably, this would limit interference with FIX onset, where blinks were linked to an increased likelihood of direction errors.

### Performance Costs of Inefficient Blink Timing

Our findings support that blink timing is highly controlled to minimize anti-saccade errors and thus maximize cognitive performance in the IPAST. We demonstrated that participants who made the most anti-saccade errors had an elevated blink probability during the appearance of critical stimuli (FIX and STIM) and decreased blink probability between trials (ITI; Fig. 4A), as well as slower learning rates and less change over the course of the task (Fig. 4C). Altogether, these findings indicate that the same deficits in inhibitory control or task engagement responsible for anti-saccade errors (Munoz & Everling, 2004) might also produce inefficient patterns of blink behavior. It is likely that inappropriate blink timing would compromise anti-saccade trials by interfering with the processing of task-relevant visual stimuli, thus compounding the downstream effects of poor inhibitory control on task performance. For example, a blink at FIX onset would delay perception of the trial instruction, shortening the time to generate the preparatory signals necessary to perform a correct anti-saccade (Coe & Munoz, 2017).

Strategically timing one’s blinks is understood as a critical part of efficient visual processing. However, more recent work suggests that blinks, timed appropriately, can also be a tool for temporarily boosting visual performance (Jit & Maus, 2020; Rolfs & Hübner, 2024; Yang et al., 2024) or providing a reset to attentional networks during a cognitive task (Nakano et al., 2013; Siegle et al., 2008). This would explain why participants do not broadly suppress blinks in the IPAST and instead, despite the cost to maintaining vision, blink rates are higher than on less cognitively demanding visual tasks (Pitigoi et al., 2024). The majority of blinks occur in the middle of the ITI, potentially representing a cognitive reset after completing a trial. The second most likely time to blink is during the middle of FIX, which could conceivably facilitate the perception and consolidation of the stimulus color and the corresponding instruction for the trial. Blinks being more strongly suppressed here on anti-saccade trials are attributed to preparatory signals related to inhibitory control (Pitigoi et al., 2024), and thus likely override any perceptual or cognitive advantage in this epoch on anti-saccade trials.

### Age-Related Changes in Optimizing Blink Timing

Despite spontaneous blinks not being consciously controlled, we infer that learning optimal blink timing requires (1) prediction of stimulus timing, (2) evaluation of the stimulus’ relative importance to the task, and (3) development of a conceptual framework for when blinking would be more or less disruptive to task performance. Blink optimization and adaptation is therefore likely to depend on high-level functions such as pattern recognition, working memory, inhibitory control, and learning capacity—cognitive domains which vary with age (Pettigrew and Martin, 2014; Clark et al., 2015; Reynolds et al., 2022; Levi and Heled, 2024). The present study is the first to examine age-related changes in blink adaptation across the lifespan.

Although neither overall blink rate nor blink duration differs by age (Pitigoi et al., 2024), we found that children and elderly participants were most likely to blink at disruptive times (during FIX and STIM onset), whereas early adults (21-24) were least likely to do so (Fig. 5). These findings support that blink patterns become more optimal throughout childhood and adolescence, then decline later in life—similar to the trajectory of other oculomotor behaviors in this task and importantly, anti-saccade error rates (Huang et al., 2024; Yep et al., 2022). Our age analysis of blink behavior is thus inextricably linked to that of task performance: age-related changes in frontal-parietal and frontal-striatal circuitry mediate changes in inhibitory control of saccade behavior across the lifespan (Ferguson et al., 2021; Luna et al., 2010). Due to the probable role of inhibitory control in blink suppression (Berman et al., 2011; Pitigoi et al., 2024) and the shared circuitry between blinks and saccades (Evinger et al., 1994), we believe these mechanisms are simultaneously driving differences in blink optimization across the lifespan.

Notably, the peak in the middle of FIX appears to be delayed in older adults relative to younger age groups (Fig. 5B). This suggests that the changes occurring during healthy aging are not simply the reverse of those occurring throughout childhood/adolescence. Possibly, they are mediated by different neural circuits (Craik and Bialystok, 2006) or reflect cognitive restructuring rather than deterioration alone (Brown et al., 2022).

With regard to the process of adaptation itself, children show less change across the task relative to both early and older adults (Fig. 5D). Yet, we caution that the steepness of the learning trajectory for early adults is likely underestimated here, because the smoothing function does not adequately capture changes in the first 10 trials, where we saw rapid changes occurring. Closer examination of this period reveals in greater detail that early adults had the same starting point as the other groups yet quickly optimized (Supplementary Figure 1). It therefore seems appropriate to conclude that early adults adapted their blinking more quickly and effectively than both children and elderly participants, consistent with the prevailing theory that learning capacity matures throughout adolescence, then declines with aging (Janacsek et al., 2012; Lukács & Kemény, 2015; Matamales et al., 2016)

### Limitations and Future Directions

The present study allows for a greater understanding of the learning process behind blink organization for optimizing goal-directed behavior. Unraveling this cognitive component—and how it varies across age and performance levels—supports that blink behavior would proxy high-level brain functions, particularly those linked to dopamine (review by Jongkees & Colzato, 2016). However, we have yet to describe blink-saccade coordination, which is necessary to provide a more complete picture of optimization within the visual system. Are there other learning processes taking place (i.e., speed of acquisition of fixation, speed of pro-/anti-saccade execution)? Are they linked to blink optimization and occur at the same rate, or are these adaptations staggered and dependent on the previous? It would also be interesting to interrogate these learning processes within a version of the IPAST in which epoch durations periodically changed—would blink timing be specifically re-learned several times? Or would a more generalizable strategy prevail?

A computational model of blinking in the IPAST would further help to identify neural pathways controlling various aspects of the behavior, and eventually to provide novel insight into the circuitry that is dysregulated in various groups. We previously showed that blinks are less efficiently structured as patients with Parkinson’s disease progress from cognitively normal to mild cognitive impairment to dementia (Brien et al., 2023). This approach would be strengthened by a better understanding of the relationship between behavioral alterations (i.e., blink patterns, adaptation) and cognitive or motor deficits localized to specific brain networks.

## Data availability

We do not have ethics board approval to make this data publicly available.

## Author contributions

ICP: Conceptualization, Formal Analysis, Visualization, Writing – Original Draft, Writing – Review & Editing

DCB: Conceptualization, Methodology, Software, Writing – Review & Editing

HCR: Investigation, Writing – Review & Editing

BKN: Investigation, Writing – Review & Editing

RY: Investigation, Writing – Review & Editing

OGC: Formal Analysis, Investigation, Writing – Review & Editing

RHK: Investigation, Writing – Review & Editing

BCC: Data Curation, Methodology, Software, Writing – Review & Editing

MM: Supervision, Writing – Review & Editing

DPM: Conceptualization, Funding Acquisition, Methodology, Supervision, Writing – Review & Editing

## Acknowledgements

We thank Drs. Gunnar Blohm and Tomas Sieger for statistical consultation in support of this project and the Munoz lab for critical feedback.

