## Supplemental Figure 1 for "Learning to blink strategically is crucial to performance in a predictable saccade task and varies across the lifespan"

**Supplemental Material**

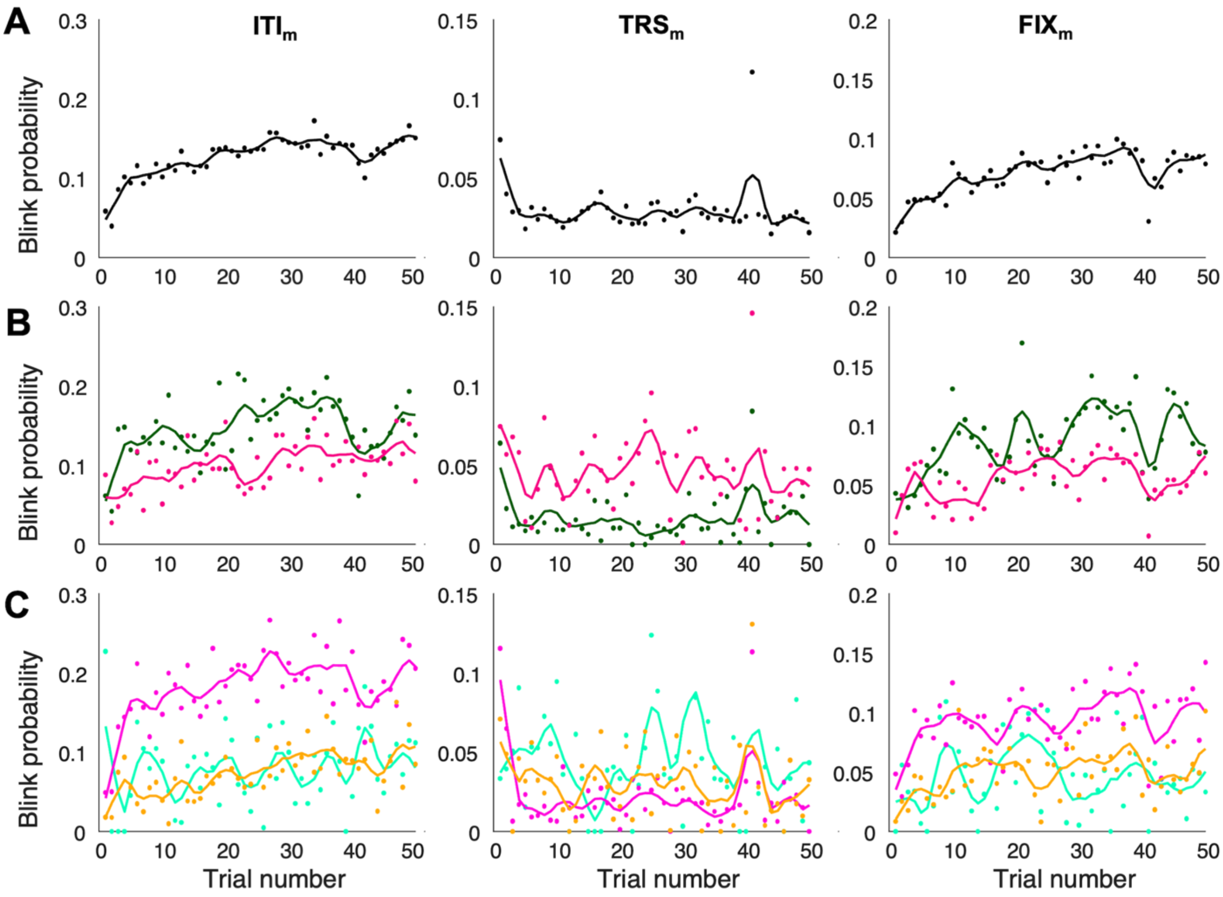


**Supplementary Figure 1. Minimally smoothed learning trajectories for (A) All participants; (B) LO and HI error groups; (C) Age groups (YC, EA, OA).** Average blink probability on every trial calculated during ITI$_m$ (far left), TRS$_m$ (middle), and FIX$_m$ (far right). Each point represents an individual trial averaged across all participants in that group. X-axis constrained to show the first 50 of 240 trials for closer examination. Minimal smoothing was applied using MATLAB smooth function with a parameter of 0.025.
